# Assessing the impact of parental linear gene normalization on the performance of statistical models for circular RNA differential expression analysis

**DOI:** 10.64898/2026.03.06.710045

**Authors:** Erda Qorri, Valentin Varga, Katalin Priskin, Dóra Latinovics, Bertalan Takács, Emese Pekker, Gábor Jaksa, Bernadett Csányi, László Torday, Ali Bassam, Zsuzsanna Kahán, Lajos Pintér, Lajos Haracska

## Abstract

**Background:** Circular RNAs (circRNAs) emerged as promising non-invasive cancer biomarkers due to their stability, abundance in body fluids, and regulatory potential. However, circRNA differential expression analysis (DEA) remains challenging, largely owing to lack of consensus on important preprocessing strategies such as filtering and normalization. While well-established bulk RNA-sequencing frameworks are commonly applied to circRNA data, newer approaches such as CIRI-DE (part of CIRI3 suite) integrate both linear and circular transcript information to improve detection. Despite developments, an assessment of these integrative strategies is lacking, and the critical impact of filtering on DEA model performance has not been comprehensively evaluated.

**Results:** In this study, we evaluated the impact of multiple normalization and filtering strategies on circRNA DEA using five experimental datasets, including two in-house blood platelet sets and semi-parametric simulated *in silico* datasets. Our results emphasize the importance of selecting an appropriate filtering threshold, as overly lenient filtering substantially reduced model performance across datasets. We found edgeR’s filterByExpr() strategy particularly effective in handling zero counts in circRNA data, while also generating the most reliable results across most datasets. Furthermore, by incorporating linear and circular information as described in CIRI-DE, most methods identified a higher number of differentially expressed (DE) circRNAs compared to circular counts alone. Notably, circRNAs identified by both CIRI-DE and the modified bulk RNA-sequencing pipelines showed substantial overlap.

**Conclusion:** Our findings demonstrate that automated filtering combined with linear-aware normalization significantly enhances the sensitivity and reproducibility of circRNA DEA, providing a standardized framework for more reliable biomarker discovery in transcriptomic research.

## 1 Background

Next-generation sequencing (NGS) has become a substantial tool in biomedical research and clinical diagnostics, enabling large-scale, high-resolution analysis of genetic material. It plays a critical role in profiling genomes and transcriptomes to link pathological phenotypes to underlying molecular changes (Weldy & Ashley, 2021). This technological leap shed new insight into the diversity of RNA molecules with direct clinical utility. Among these, circular RNAs (circRNAs) have become key players in cancer and other diseases, as NGS has enabled their identification, quantification, and functional characterization (Byron et al., 2016). However, effective non-invasive detection of cancer remains a significant challenge primarily due to the scarcity of stable biomarkers and technical difficulties in their precise quantification. This issue is especially critical in liquid biopsy, where tumor-derived nucleic acids are present at very low concentrations and rapidly degrade within bodily fluids (Best et al., 2018; Heitzer et al., 2019). However, the use of platelets in blood samples presents a promising approach to address the problem of low RNA abundance, being enriched in linear and circular RNAs at significantly higher rates than in other cell types (Preußer et al., 2018).

circRNAs are a special class of RNA molecules that are formed when the linear splicing of pre-mRNA is replaced by a back-splicing event, resulting in a covalently closed, continuous loop. This unique topology provides remarkable resistance to exonuclease degradation and also separates circRNA from its linear counterparts, enabling independent regulation and function within the cell. circRNAs are not merely byproducts but are selectively produced in a cell-type-, tissue-, and developmental stage-specific manner and are often conserved across species, indicating evolutionary significance (Leypold & Speicher, 2021).

Functionally, circRNAs serve as versatile regulators within the cells. Their exceptional stability, abundance in body fluids and extracellular vesicles further emphasize their potential as robust biomarkers for disease diagnostics and cancer monitoring. Emerging evidence supports that platelet-derived circRNAs can mirror early tumorigenic events, making them potentially ideal for non-invasive cancer biomarkers. Crucially, given the subtle expression changes characteristic of early-stage cancer, highly reliable statistical models are essential to accurately distinguish biological signal from noise and optimize diagnostic accuracy.

In RNA-based molecular analyses, accurate detection of changes in transcript abundance is fundamentally dependent on precise relative quantification and rigorous definition of parameters and standards throughout the experimental and analytical workflow. Scientific consensus underscores that the reliability of DEA hinges on the minimization of technical and biological biases and the careful selection and validation of normalization strategies and reference standards. While NGS technologies have matured, unbiased and accurate RNA quantification remains a moving target, with normalization standing as a central, evolving challenge for robust transcriptomic analysis (Conesa et al., 2016).

The emergence of circRNA as a transcriptomic feature has introduced a new layer of bioinformatic difficulty, as standard linear RNA analysis tools are insufficient, and specialized algorithms are required to accurately identify back-splice junctions (BSJs) unique to circRNAs (Digby et al., 2024; Drula et al., 2024).

Currently, numerous tools are available for detecting circRNAs from total or Ribo-depleted RNA libraries (Westholm et al., 2014; Wang et al., 2010; Ma et al., 2021; Zheng et al., 2025). These tools primarily rely on the identification of chimeric reads corresponding to BSJs, a hallmark characteristic of circRNAs (Nielsen et al., 2022). However, no widely accepted standard exists regarding the most reliable approach for circRNA identification and filtering of the obtained results. Benchmarking studies on both short- and long-read sequencing demonstrate that present algorithms identify different numbers of circRNAs and are susceptible to false positives. To achieve more reliable results, a concordance-based strategy combining the outputs of multiple bioinformatics tools has become widely accepted practice in the field. In fact, Vromman et al., 2021 demonstrated an increase in detection sensitivity when several algorithms were utilized in combination. While computational tools for circRNA detection have matured, consensus on the best practices for identifying DE circRNAs across biological conditions is lacking. DEA methods originally developed for bulk RNA sequencing, such as DESeq2, edgeR, and limma-voom, remain widely utilized, despite recent findings demonstrating that circRNA BSJ counts follow distributions more typical of single-cell RNA sequencing or metagenomic data than linear RNA, indicating that conventional DEA models are often unable to fully account for the sparse BSJ counts present in circRNA data. Moreover, recent studies have shown that use of default parameters can yield suboptimal and potentially misleading results, emphasizing the need for careful parameter tuning (Maza et al., 2013).

Another important consideration in circRNA DEA is the normalization strategy, which is typically based on BSJ-spanning reads, thus directly taking the abundance of circular counts as the expression level of circRNAs. However, this approach can be problematic as it overlooks the unique expression dynamics of circRNAs relative to their host genes as well as evidence suggesting that circRNA can be regulated independently of its parental linear RNA (Alonso-García et al., 2025).

To address these challenges, new approaches tailored that account for the unique properties of circRNAs. These methods often incorporate linear RNA expressions when estimating circRNA abundance (L. Chen et al., 2020; Zhang et al., 2020). Notably, CIRI3 utilizes the normalization strategy described in CIRIquant, in which normalization factors derived from linear RNA are subsequently applied to normalized circRNA expression (Zheng et al., 2025). However, the effectiveness of these novel normalization methods has not been evaluated in comprehensive benchmarks, representing a key gap in the field.

To address these issues, in this study, we performed a comprehensive benchmarking of commonly utilized DEA methods on a combination of both simulated and experimental data derived from non-enriched total RNA-sequencing datasets, including two in-house platelet datasets from early breast cancer patients. In addition, we systematically characterized the sparse and zero-inflated nature of circRNA counts and explored the impact of three filtering strategies on the performance of widely utilized differential expression algorithms. Furthermore, through a structured three-step workflow, we explored the benefits of incorporating information derived from linear RNAs into circRNA DEA. Collectively, our findings highlight the limitations of treating circRNAs in isolation from their linear counterparts and support the development of circRNA DEA methods tailored to their unique nature.

## 2 Results

### 2.1 Characteristics of circRNAs in different non-circRNA-enriched datasets

To conduct this study, we employed a combination of publicly available datasets obtained from the Sequence Read Archive (SRA) and simulated datasets, along with our own liquid biopsy data. Public datasets were selected from peer-reviewed studies according to the following previously described criteria: (i) a minimum of four biological replicates per condition, (ii) a sequencing depth of at least 20 million reads per sample, and (iii) clear separation between the samples within each condition (Buratin et al., 2023) (Figure 1). An additional key selection criterion was the library preparation strategy; we specifically included datasets generated from total RNA libraries.

**Figure 1.**
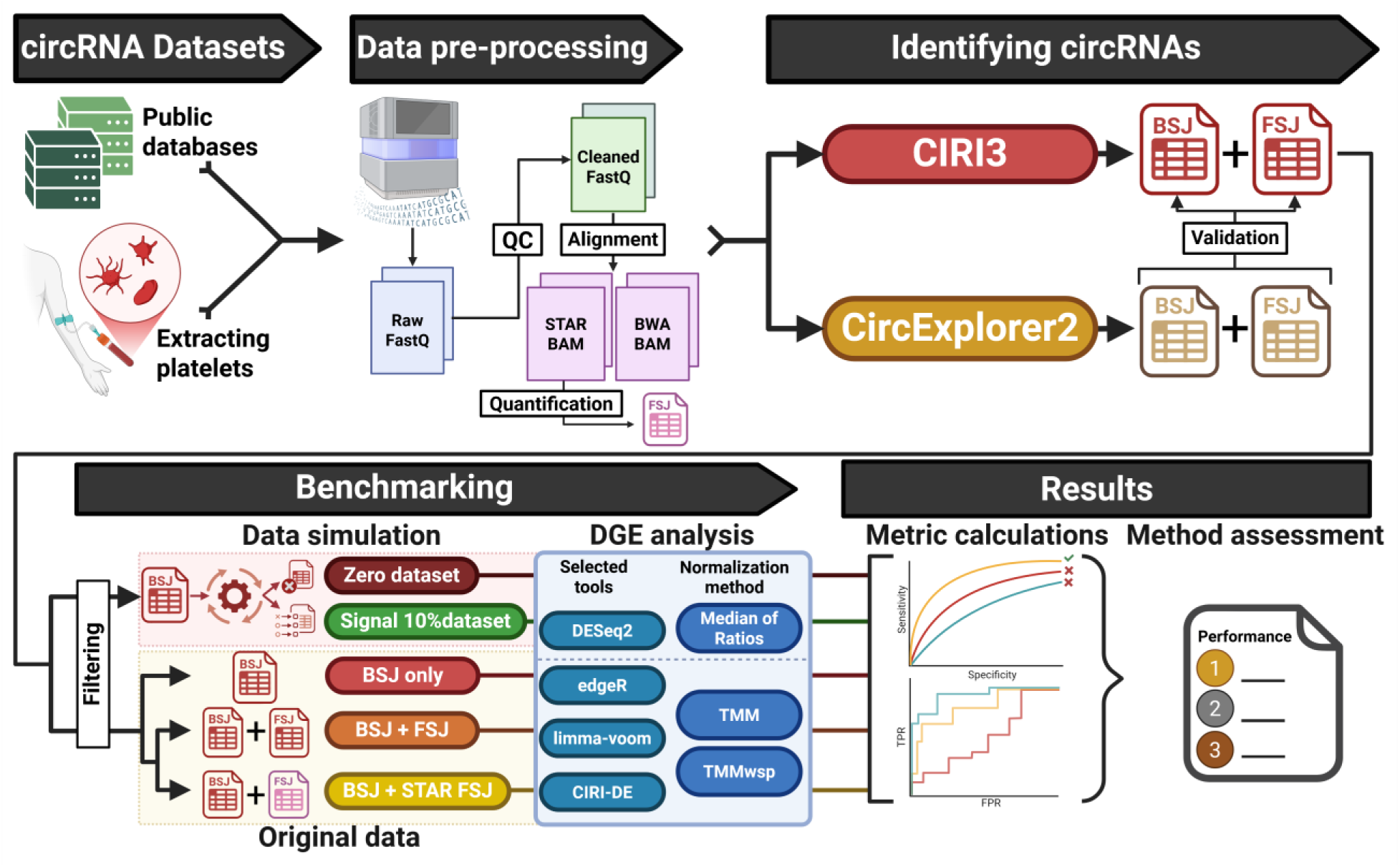
Workflow of the benchmarking steps starting with dataset acquisition, data processing, simulation, and performance evaluation. Image was generated using BioRender.

Emphasis was placed on total RNA libraries, since the majority of recent circRNA quantification studies reported in the literature rely on total RNA isolation rather than circRNA-specific enrichment (L.-L. Chen, 2020). Based on the abovementioned criteria, three public datasets with a total of 84 samples were selected (Table 1; Supplementary Figure 1).

**Table 1.**
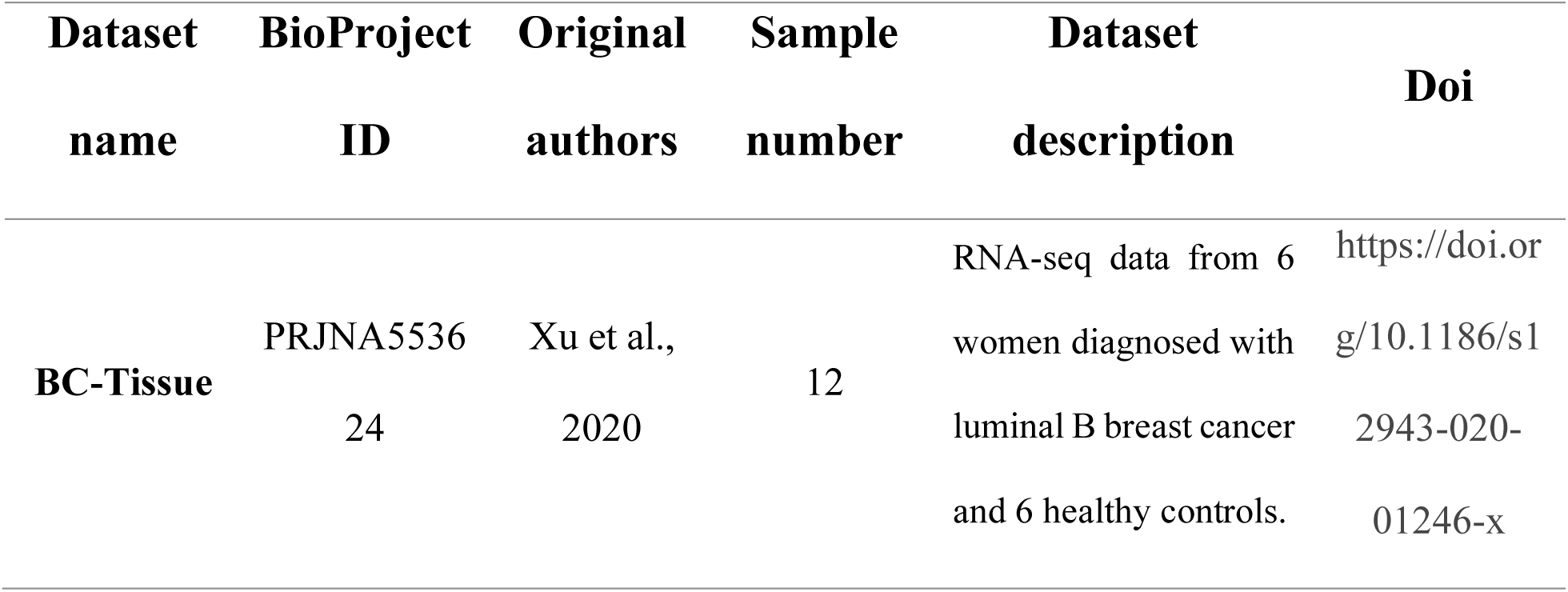
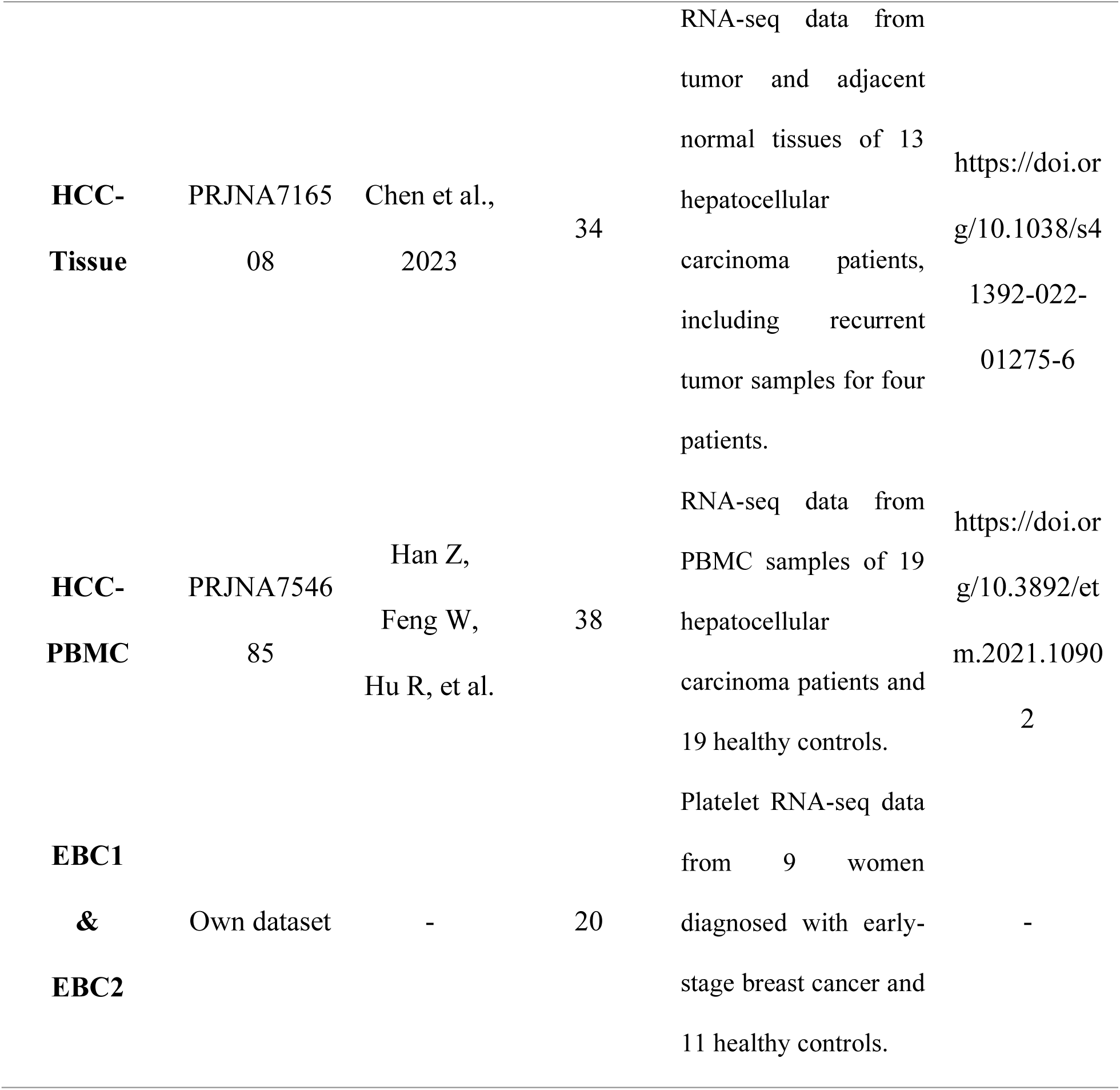
Selected datasets. BC: Breast Cancer; HCC: Hepatocellular Carcinoma; PBMC: Peripheral Blood Mononuclear Cell; EBC: Early-stage Breast Cancer For all datasets, total RNA was subjected to rRNA depletion, converted to cDNA, and sequenced using paired-end Illumina technology (2 × 150 bp). Libraries were strand-specific where applicable.

Simulated datasets were generated following previously reported strategies described by Buratin et al. (2023), as detailed in the Methods Section. In addition, we analyzed an in-house dataset consisting of blood platelets isolated from healthy individuals and patients with early-stage breast cancer (EBC). Owing to the established high circular-to-linear RNA ratio in platelets, this dataset was considered a natural model of circRNA-enriched libraries, providing a unique source for the benchmark.

For the identification of BSJ and forward-spliced junction (FSJ) events across all datasets, we employed two widely utilized circRNA-detection algorithms: CIRI3 (Zheng et al., 2025) and CircExplorer2 (Ma et al., 2021). The use of multiple algorithms was intended to minimize algorithm-specific biases in circRNA detection and to increase the reliability of circRNA identification, as recommended by previous studies (Hansen et al., 2016; López-Jiménez & Andrés-León, 2021; Vromman et al., 2023). Although there are currently no standard guidelines regarding the optimal number or specific combination of circRNA-detection algorithms, our selection was guided by their widespread use in the field and reported performance in literature.

Our results showed that the number of identified circRNAs varied substantially across datasets (Figure 2A), which can be attributed to the differences in both sample size and sequencing depth. We observed that CE2 identified a higher number of circRNA candidates than CIRI3 in most datasets, except for the HCC-PBMC dataset. However, after the filtering steps, the number of confidently identified circRNAs by the two algorithms converged to a comparable range (Figure 2A). To ensure that both CIRI3 and CE2 identified the same circRNAs with high confidence, we assessed the concordance by comparing circRNA identifiers. This analysis showed a high overlap of the retained circRNAs, which was also supported by high Jaccard similarity indices (0.89 - 0.99; Figure 2B).

**Figure 2.**
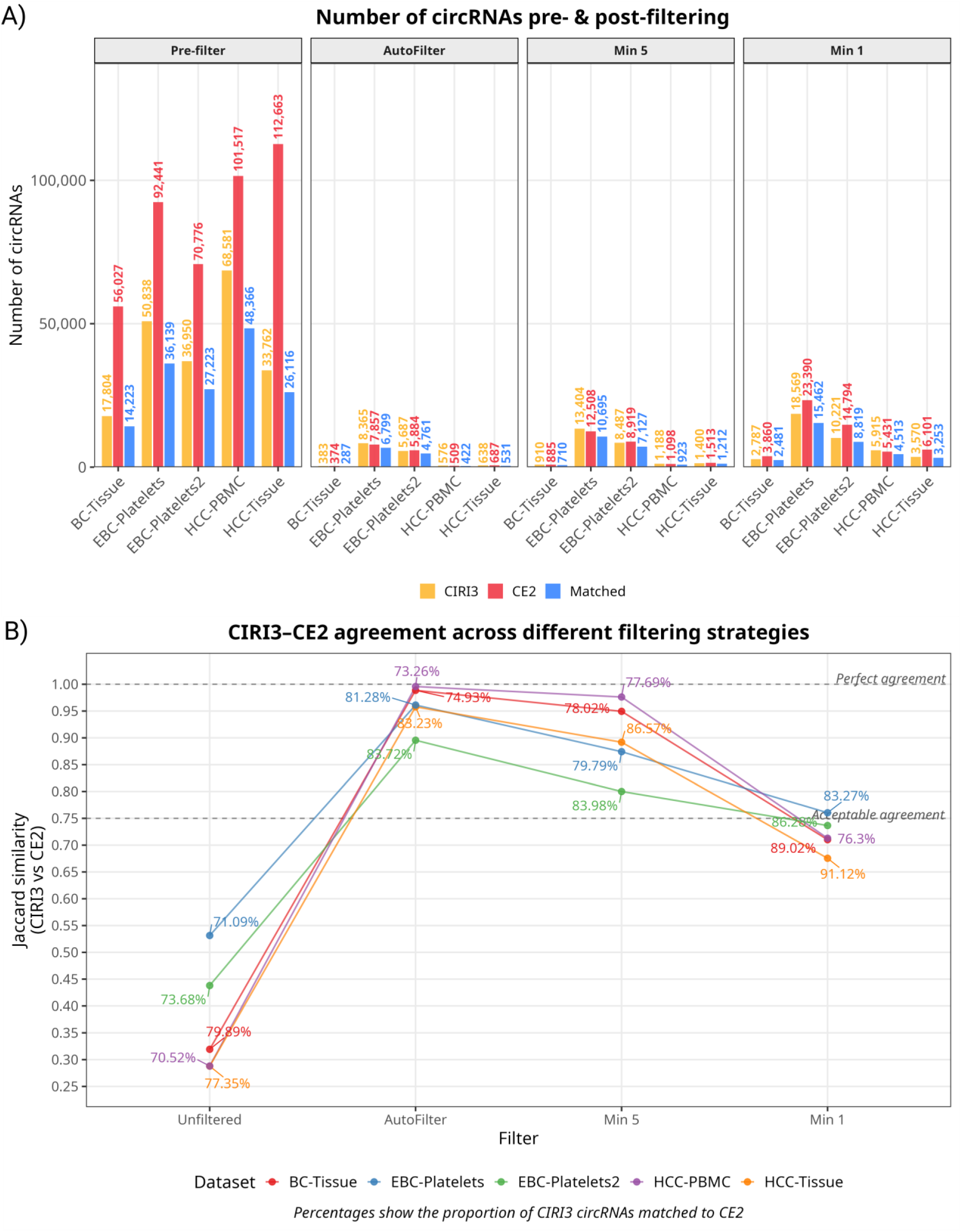
circRNA identification. (A) Total number of circRNAs identified with CIRI3 and CE2 before and after applying automated filtering across all datasets. (B) Jaccard similarity index between CIRI3 and CE2 across the different filtering strategies. Each line represents a dataset; points indicate the similarity index value at each filtering threshold. Percentages show the proportion of CIRI3-detected circRNAs that were successfully matched to a CE2 counterpart. BC: Breast Cancer; HCC: Hepatocellular Carcinoma; PBMC: Peripheral Blood Mononuclear Cell; EBC: Early-stage Breast Cancer.

Next, we evaluated the percentage of zero counts of the identified circRNA in each dataset. The results obtained showed that the percentage of zero counts was high across all datasets, independent of the library preparation strategy. Notably, the first platelet dataset exhibited lower proportions of zero counts compared to the other datasets, with values reaching 63.05 % and 67.59 % for CIRI3 and CE2, respectively (Table 2).

**Table 2.**
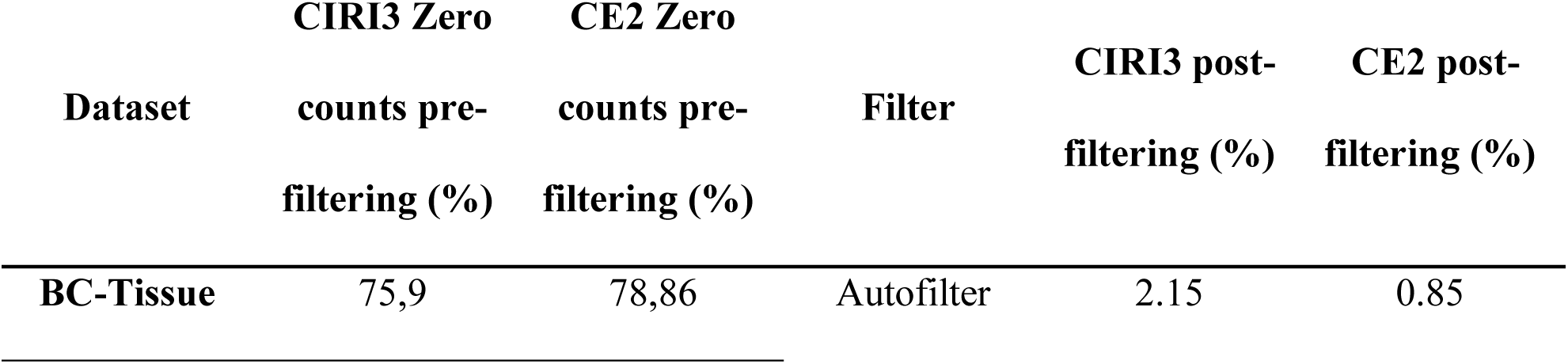
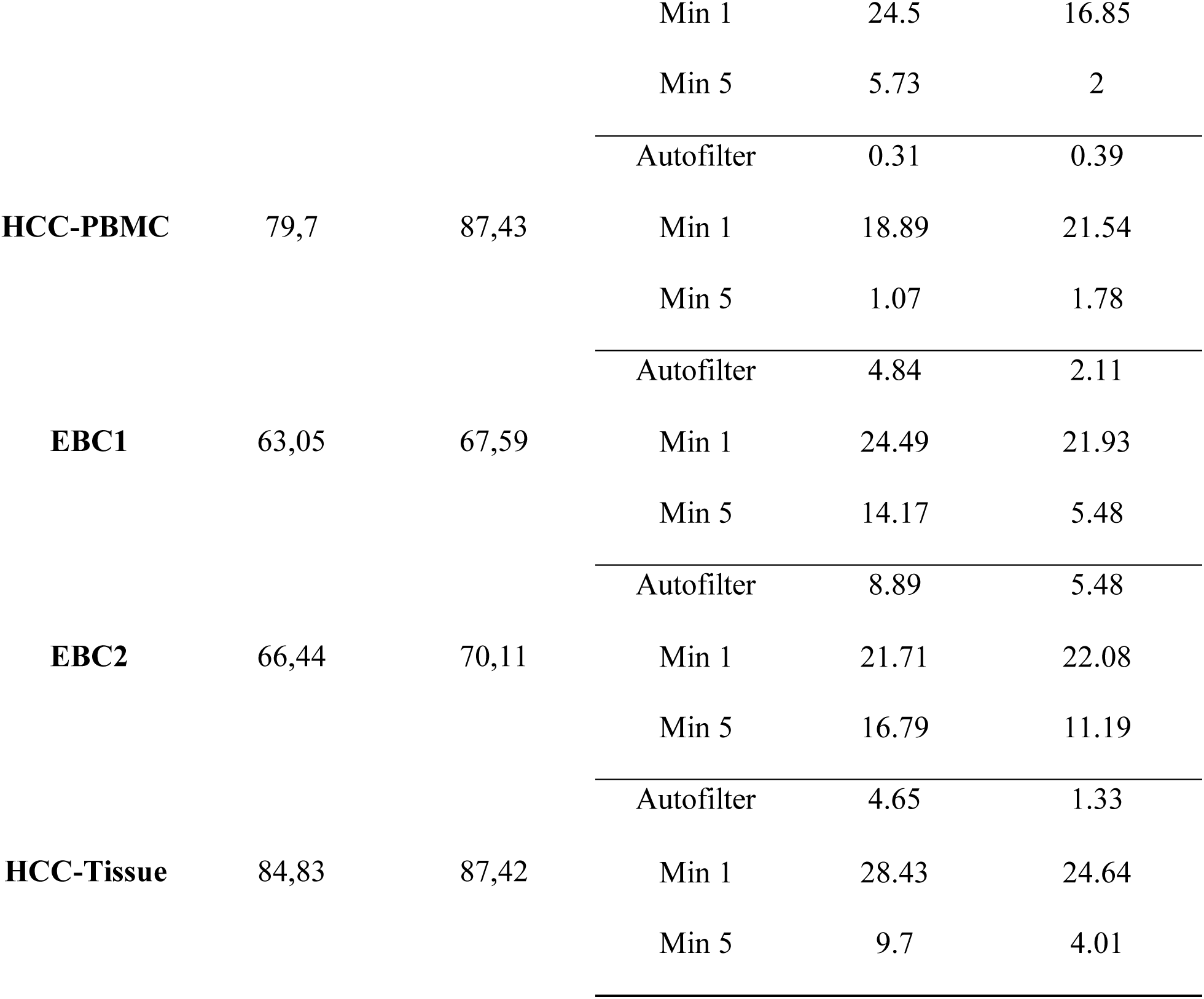
Percentage of circRNAs with zero counts prior to and after different filtering strategies, for both CIRI3 and CircExplorer2 (CE2). BC: Breast Cancer; HCC: Hepatocellular Carcinoma; PBMC: Peripheral Blood Mononuclear Cell; EBC: Early-stage Breast Cancer

To address the zero-inflated nature of circRNA datasets, several filtering strategies have been proposed; however, due to the lack of standardization in the literature and the potential bias introduced by arbitrarily defined thresholds, we adopted the automated filtering approach implemented in edgeR’s filterByExpr(), which we refer to as auto-filtering from this point onwards. As expected, this filtering step substantially reduced the proportion of zero counts across all datasets, ranging from 0.31% in the HCC-PBMC dataset to 6.57% in the platelet dataset as reported by CIRI3 (Table 2). By contrast, testing less stringent filtering (Min 5 and Min 1) resulted in a higher proportion of zero counts and low-abundance circRNAs retained compared to auto-filtering. Specifically, under Min 5 filtering, zero counts retained ranged from 1.07% (HCC-PBMC) to 16.79% (EBC2) for CIRI3, and from 1.78% to 11.19% for CE2. This effect was substantially enhanced under Min 1 filtering where retained zero counts increased from 18.89% to 28.43% for CIRI3, and from 16.85% to 24.64% for CE2 across datasets.

The impact of selecting an adequate filtering strategy was particularly evident in the estimation of Jaccard similarity across the datasets (Figure 2B). Without filtering, pairwise Jaccard similarity values ranged from 0.29 to 0.53 across datasets. However, following auto-filtering, these values increased notably, ranging from 0.68 to 1.00, suggesting that removing low-abundance circRNAs reduces stochastic variability and increases inter-method concordance.

Consistent results were also obtained using CE2, further indicating that the observed patterns reflect intrinsic properties of circRNA expression in the datasets rather than algorithm-specific detection biases (Figure 2B).

As circRNA identification primarily relies on the detection of BSJs, we next examined the distribution of read counts supporting such events across all datasets. Prior to filtering, the median BSJ read count was zero across all datasets, as expected given the high proportion of zero counts, while under auto-filtering, the median ranged from 16 to 25 read counts per sample. Interestingly, median BSJ read counts dropped consistently when applying Min 5 (9-15.5 reads per sample across datasets) and Min 1 filtering (2-11.5 reads per sample), reflecting the inclusion of many poorly supported circRNAs. These results were consistent between CIRI3 and CE2 across all filtering strategies, highlighting the influence of filtering choice on overall BSJ support levels. Moreover, after filtering, the difference between CIRI3 and CE2 was remarkably reduced (Figure 3A). Notably, the circRNA-enriched platelet libraries (EBC1 and EBC2) exhibited higher median BSJ read counts compared to the Tissue and PBMC datasets across all filtering strategies, with median counts of 20.5-25.5 reads per sample under default filtering compared to 15.5-17.5 in the remaining datasets, consistent with the known enrichment of circRNAs in platelets. This enrichment advantage was most apparent under auto-filtering, where platelet datasets showed median BSJ counts 5-8 reads higher than the Tissue and PBMC datasets. While the absolute difference narrowed under more stringent filtering (Min5: 2-5 reads and Min 1: 3-7 reads), the platelet datasets consistently retained the highest median BSJ support across all thresholds, suggesting that the circRNA enrichment in platelets is robust to the filtering strategy.

**Figure 3.**
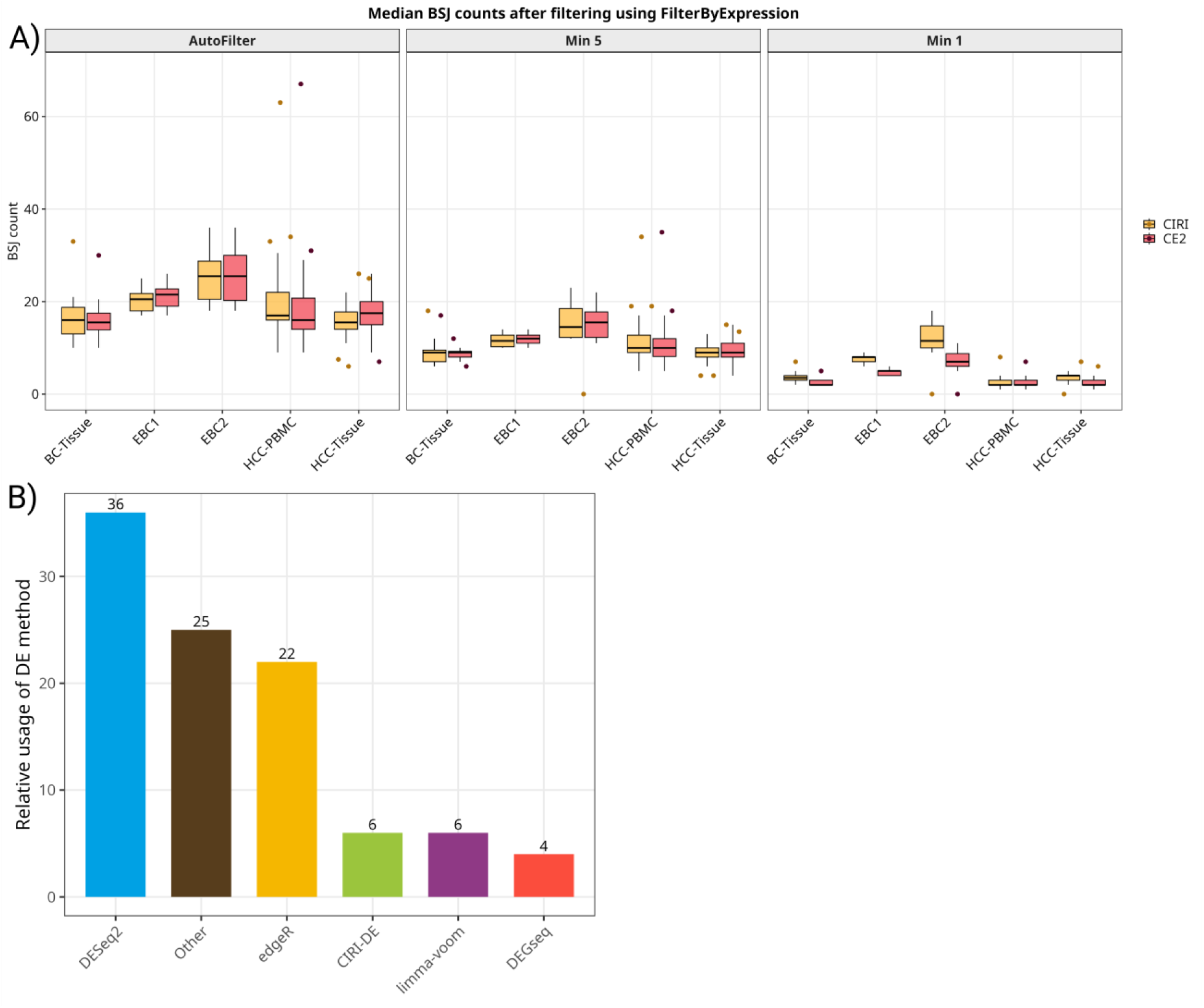
Boxplots summarize sequencing depth across samples; n indicates the total number of samples per dataset. (A) Median number of reads supporting BSJ events detected by CIRI3 and CircExplorer2 (CE2) across datasets and filtering strategies. (B) Relative usage of differential expression analysis tools for circRNA studies in the literature. BC: Breast Cancer; HCC: Hepatocellular Carcinoma; PBMC: Peripheral Blood Mononuclear Cell; EBC: Early-stage Breast Cancer

To ensure that differences between conditions in the selected datasets were sufficient to support circRNA detection, we performed unsupervised principal component analysis (PCA) using BSJ and FSJ counts separately, as well as in a combination. The combined use of BSJ and FSJ counts was intended to reflect a novel normalization approach implemented in CIRI3 (Zheng et al., 2025). To further evaluate whether linear RNA information could be leveraged to normalize and model circRNA expression, PCA was also performed on total RNA libraries normalized using normalization factors derived from linear read counts.

The resulting PCA analyses showed tight within-condition clustering, supported by a moderate-to-high inter-sample correlation for each condition (Supplementary Figure 2), together with clear separation between conditions across all datasets, regardless of the origin of the read counts. These results indicate that the transcriptional differences between the conditions appear to be consistently captured by both circular and linear RNA signals, making them appropriate to model DEA. A comparable separation between conditions was observed, validating the FSJ-based analysis (Supplementary Figure 2). The BSJ log2CPM densities across the different normalizations are shown in Supplementary Figure 3.

### 2.2 DEA method selection

Given the high proportion of zero counts observed across the datasets and the notable impact of filtering strategies in reducing them, we sought to evaluate how different filtering thresholds influence the behavior of differential expression models in identifying DE circRNAs. In the absence of standardized guidelines for selecting DEA tools for circRNA data, we selected methods based on their usage frequency in the field as defined by the number of recent citations. As expected, most research carried out in the field relies on three well-established bulk RNA-sequencing pipelines, DESeq2, edgeR, and limma-voom (Figure 3B).

A substantial number of studies implement custom DEA strategies or combinations of already existing pipelines, further emphasizing the lack of uniformity in the field. Approaches like CIRI-DE which integrate both linear and circRNA-derived information prior to DEA are among the least frequently utilized methods.

Given the absence of clear filtering guidelines, we assessed the performance of DESeq2, edgeR, and limma-voom under different filtering thresholds and parameters, along with CIRI-DE’s linear RNA-aware framework to determine whether incorporation of this linear information improves circRNA DEA detection over the current bulk RNA-sequencing methods.

To evaluate their performance, we simulated 30 and 50 replicates for the BC, EBC1, and EBC2 datasets and the HCC-PBMC and HCC-Tissue datasets, respectively. Among the simulated datasets, one was a “Zero Set” dataset (no DE circRNAs) and a DE-Signal-10% dataset (10% DE circRNAs). In total, 1140 individual datasets were utilized subsequently as input for downstream analyses.

### 2.3 Type 1 error control

The performance of the selected DEA methods in controlling Type 1 error was evaluated across five simulated datasets that did not contain any DE circRNAs. circRNAs incorrectly identified as DE were automatically considered false positives and used to calculate the false-positive rate (FPR) for each method. FPR values were calculated at three thresholds (0.01, 0.05, and 0.10) applied to the unadjusted p-values reported by the DEA tools.

Next, given the zero-inflated nature of circRNA data, we examined the impact of different filtering strategies on FPR values across all tested DEA tools and datasets. Notably, the voomLmFit pipelines using both Trimmed Mean of M-values (TMM) and TMMwsp normalization consistently displayed inflated FPR values across all datasets and filtering thresholds, especially under Min 1 filtering, reaching 0.18 and 0.15 in the BC dataset, respectively (Supplementary Tables 1 and 2). The remaining limma-voom implementations displayed the opposite behavior, with median FPR values falling generally below the nominal level, indicating a tendency toward conservative Type I error control. This behavior was more pronounced under the two lenient filtering thresholds, Min 1 and Min 5, and was particularly consistent in the two platelet datasets, EBC1 and EBC2.

DESeq2 configurations showed minimal sensitivity to the applied filtering thresholds, with all three-producing similar, near-nominal results in the case of auto-filtering and Min 5. However, Min 1 filtering results indicate a conservative Type 1 error rate control, characterized by FPR values consistently below the nominal 0.05. Similar results were observed for edgeR, with TMMwsp normalization reducing the median FPR close to nominal under auto-filtering as opposed to TMM.

Overall, auto-filtering provided the most balanced performance across the tested methods, with FPR values close to the nominal 0.05 level and comparatively low inter-dataset variability. Min 5 filtering produced similar results, although it was characterized by greater dispersion in certain datasets (Figure 4A).

**Figure 4.**
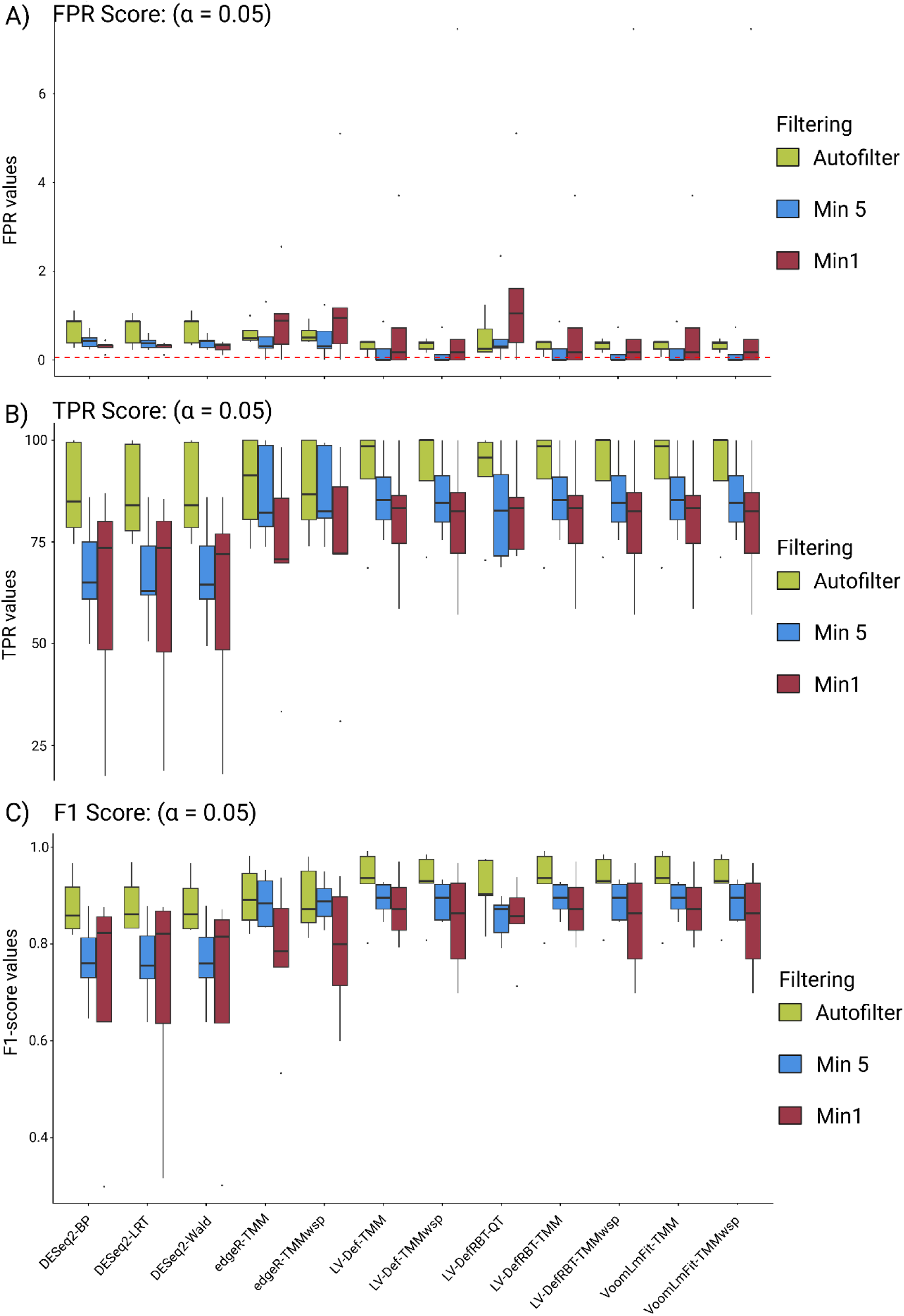
Performance of the differential expression analysis method and their different parameter combinations across all DE-Signal datasets and filtering strategies. The boxplots show (A) the distribution of median false-positive rate, (B) true-positive rate (sensitivity), and (C) F1 scores across the different datasets. These results are reported at an alpha score of 0.05.

### 2.4 Performance evaluation on the DE-Signal-10 dataset: FPR, TPR, F1 score and area under the precision–recall curve (AUPRC)

TPR varied notably across filtering thresholds for all evaluated DEA methods. Auto-filtering consistently resulted in the highest sensitivity with median TPR values across all datasets ranging from 0.84 in DESeq2-Wald to a perfect score in the three limma-voom pipelines, LV-Def-TMMwsp, LV-DefRBT-TMMwsp, and LV-voomLmFit-TMMwsp.

By contrast, under Min 1 filtering, median TPR values of all methods decreased considerably, accompanied by an increase in inter-dataset variability. This effect was particularly pronounced for all three DESeq2 implementations and edgeR (TMM and TMMwsp,) with median TPR values ranging from 0.707 in edgeR_TMM to 0.735 in DESeq2-BP (Figure 4B). Interestingly, DESeq2 implementations were most sensitive to the applied filtering strategy exhibiting TPR values well below 0.5 for certain datasets under Min 1 filtering but a more dataset-dependent performance in Min 5. Their performance substantially improved under auto-filtering; however, inter-dataset variability remained high as opposed to the other methods.

Limma-based implementations, on the other hand, exhibited an overall stable performance under auto-filtering, characterized by high-median sensitivity values. Across individual datasets, all methods demonstrated reduced sensitivity and increased variability in performance in the HCC-PBMC dataset and the other two liquid-derived datasets (EBC1 and EBC2), especially under Min 1 and Min 5 filtering (Figure 4A), highlighting the influence of biological input origin on method performance (Luo et al., 2020). Overall, our findings indicate that reducing filtering stringency results in increased performance variability and compromises sensitivity for several methods, particularly for DESeq2 and edgeR. In contrast, auto-filtering emerged as a plausible method which maximizes sensitivity, while stabilizing performance across different datasets.

A reliable DEA method, however, should enable the robust identification of truly expressed circRNAs, while simultaneously limiting false-positive calls. To evaluate their ability to reduce false positives, we calculated the median false discovery rate (FDR) values across all datasets for each DEA tool. The obtained FDR values varied considerably across datasets, with the impact of the filtering strategy differing depending on both the method utilized and the dataset (Supplementary Figure 4). Among the tested methods, DESeq2 displayed the highest increase in FDR under the auto-filtering, driven by elevated false-positive calls across all datasets except for EBC2. Comparably high FDR values were obtained under Min 5, while Min 1 filtering generally resulted in lower and stabilized FDR values, ranging from 0.298 in the BC dataset to FDR >0.5 in the HCC datasets. The opposite pattern held true for both edgeR and the limma-voom approaches, where Min 1 filtering led to the identification of a higher number of false-positive circRNAs and consequently higher FDR values across datasets, whereas all the tested limma-voom configurations demonstrated lower FDR values relative to both DESeq2 and edgeR. However, their performance appeared more sensitive to lenient filtering, as reflected by the increase in FDR values under Min1 filtering (Supplementary Tables 3 and 4). Although method performance was predominantly dataset dependent, auto-filtering generally resulted in lower median FDR values and reduced inter-dataset variability compared to both Min 1 and Min 5. If considered independently, sensitivity and FDR do not fully capture the ability of DEA methods to balance false-positive and false-negative instances; therefore, to obtain a more robust evaluation of their performance we calculated the F1 scores across all datasets and filtering thresholds.

All methods achieved higher F1 score values under the auto-filtering strategy, ranging from ∼0.82 for the DESeq2 versions to 0.99 across the different limma-based configurations, indicating consistently strong performance regardless of the parameter changes, when applied to circRNA data (Figure 4C). The superiority of the auto-filtering strategy was largely consistent across datasets, resulting in overall higher F1 scores compared to both Min 1 and Min 5 filtering. Under Min 5 filtering, F1 scores showed a modest decline, from 0.64 in DESeq2-LRT to 0.95 in edgeR-TMM, whereas Min 1 filtering resulted in a stronger decrease in performance, with median F1 values ranging from 0.29 in DESeq2-LRT to 0.97 in LV-Def-TMM, Limma-voom_TMM, and VoomLmFit-TMM. Interestingly, all DEA methods performed poorly in the HCC-PBMC dataset, where most achieved F1 scores >0.80 under auto-filtering but dropped considerably <80 under the less stringent filtering strategies (Supplementary Figure 5).

Notably, DESeq2 and edgeR appeared more sensitive to lenient filtering approaches, particularly for Min 1 in the case of edgeR, whereas limma-voom configurations maintained a relatively stable distribution of F1 scores across datasets, even under Min 1 and Min 5 filtering. Overall, our findings suggest that limma-voom versions provide a more balanced and stable trade-off between sensitivity and precision across different filtering conditions and heterogeneous circRNA datasets stemming from different biological sources, such as liquid biopsies and tissue.

Furthermore, evaluation of AUPRC performance showed that differential expression methods displayed better results under auto-filtering, with AUPRC values ranging from 0.89 to 1.0. In comparison, their performance slightly decreased under Min 5 (0.84–0.99) and Min 1 (0.62–1.0) filtering thresholds (Supplementary Figure 6A). Notably, Limma-voom pipelines maintained the most stable and highest scores across all datasets, including HCC-PBMC and HCC-Tissue (AUPRC > 0.96), with LV-DefRBT, LV-DefRBT-QT, and LV-Def-TMM configurations (in both TMM and TMMwsp modes) achieving an AUPRC of 1.0. Overall, these tools demonstrated consistent performance regardless of the applied filtering threshold.

A similar performance was observed for the two edgeR pipelines, which displayed only modest sensitivity to the filtering strategy, as reflected by their median AUPRC values >0.90 for both TMM and TMMwsp normalizations. However, their inter-dataset variability notably increased compared to the limma-voom methods, largely driven by a decrease in performance in both HCC datasets (Supplementary Figure 6B). Based on the obtained AUPRC values, our findings suggest that limma-voom configurations and edgeR can discriminate true positives and false positives more reliability across diverse biological contexts. Furthermore, they appeared to be less sensitive to the filtering strategy, indicating less impact on the results obtained when different approaches are applied.

### 2.5 Reproducibility and tool ranking on the DE-Signal-10 dataset

To determine whether the sets of significantly expressed circRNAs reported by the methods under an FDR threshold of 0.05 were consistent across all replicates per sample, we calculated the average Jaccard similarity index per dataset per method across the tested filtering levels. The results obtained show that none of the methods achieved a Jaccard score exceeding 0.80 in any of the dataset under any filtering strategy (Supplementary Figure 7). The highest similarity (0.758) was identified in the EBC2 datasets reported by the three Limma-voom-LmFit-TMM, limma-voom-default-TMM, and limma-DFRBT-TMM pipelines. Consistent with our previously described results, all methods displayed poor similarity scores in both HCC datasets (below 50), especially under Min 1 and Min 5 filtering. Interestingly, across the BC, HCC-PBMC, and HCC-Tissue datasets, the three DESeq2 pipelines displayed the highest consistency in the predicted circRNA sets. When evaluating the impact of filtering on their performance, all methods displayed higher similarity indices under the auto-filtering approach as compared to Min 1 and Min 5 strategies, suggesting that removing low-count circRNAs prior to DEA removes background noise, leading to more reproducible and consistent DEA calls across replicates.

### 2.6 Analyzing the effect of linear normalization on circRNA expression

To evaluate whether incorporating linear RNA information improves the detection of DE circRNAs compared to existing BSJ-only pipelines, we performed a second benchmarking analysis using five real-world experimental datasets (Table 1). The selected DEA methods, along with CIRI-DE, which implements a linear RNA-aware normalization strategy of the circRNA counts, were assessed under three scenarios. In brief, (I) the conventional BSJ-only approach, (II) the BSJ + FSJ strategy, and (III) a full linear RNA approach, in which information from the total linear RNA libraries was incorporated.

Although the performance of the DEA tools varied across datasets and significance thresholds, the obtained results consistently showed that most methods reported a few or no DE circRNA under the BSJ-only strategy, particularly at α = 0.01, where nearly all methods reported either zero or less than 10 DE circRNAs, with DESeq2 in Wald Test mode being the only exception identifying more than 13 DE circRNAs in the HCC-Tissue dataset. When incorporating information from the FSJ sites or total linear RNA, the number of reported DE circRNAs increased across the datasets (Supplementary Figures 8 to 11).

This was especially notable for CIRI-DE and the different limma-voom pipelines, which consistently identified a larger number of DE circRNAs, as compared to zero under the BSJ-only scenario. This trend became more pronounced as we applied more lenient significance thresholds such as 0.05 and 0.10, where most methods remained largely conservative in the BSJ-only scenario as opposed to the linear-aware strategies, which consistently reported more DE circRNAs across most datasets, especially in EBC1 and EBC2 (Figures 5A and 5B). Notably, CIRI-DE in both modes, FSJ and linear RNA information, reported >70 DE circRNA in the HCC-Tissue dataset. However, the response to linear-aware normalization was not uniform across all methods, as edgeR-glmQLFit (in both TMM and TMMwsp mode) and DESeq2 (BetaPrior and LRT) displayed a more conservative behavior than the limma-voom pipelines and especially CIRI-DE, as the latter consistently reported higher numbers of DE circRNAs across all datasets. Moreover, when incorporating FSJ-derived information, CIRI-DE (TMM-normalized) identified approximately two times more differentially expressed circRNAs as opposed to edgeR-QLFit TMM workflow. A similar trend was observed when CIRI-DE utilized information from the linear RNA as well.

**Figure 5.**
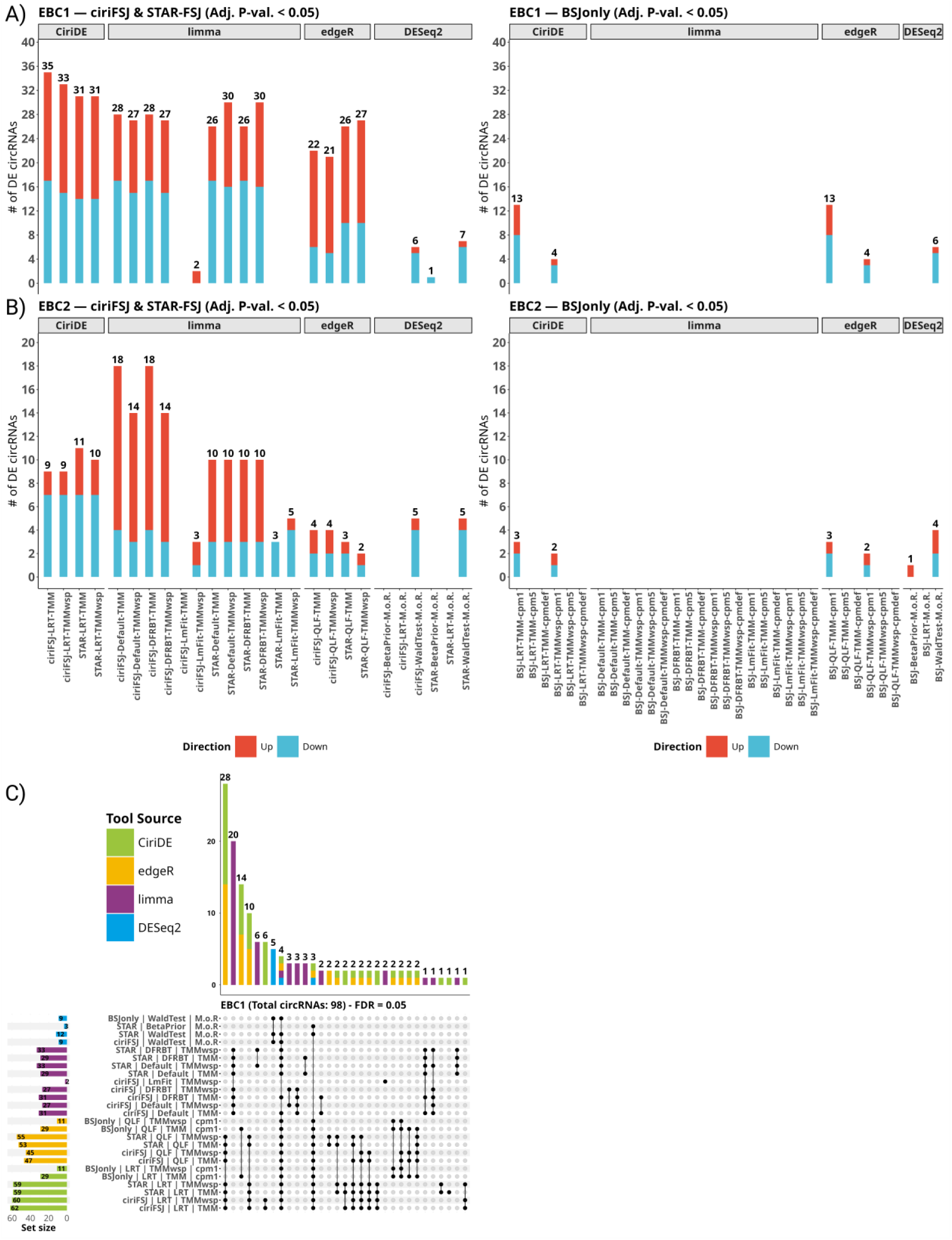
Comparison of DEA methods for circRNA detection. (A-B) Stacked barplots showing the number of significantly DE circRNAs in the EBC1 and EBC2 datasets. (C) UpSet plots illustrating the overlap of DE circRNAs across different tool-normalization combinations in EBC1 datasets.

Lastly, except for CIRI-DE and edgeR-glmQLFit identifying only one DE circRNA under the BSJ-only scenario, none of the methods reported DE circRNAs in the HCC-PBMC dataset, suggesting high within-group variance and minimal differences reflected at the circRNA level across the tested biological conditions of this dataset. These results are consistent with the findings from the simulated datasets, where most methods consistently displayed poor performance in the HCC-PBMC dataset.

Subsequently, having characterized the impact of linear-derived information between the DEA methods, we assessed whether the incorporation of this information had an impact on the directionality of expression of the identified circRNAs.

To this end, circRNAs identified across both scenarios were collected, and the consistency of their log fold changes (logFC) were evaluated using CIRI-DE’s predictions as a reference baseline. The results demonstrated that although the magnitude of the fold changes per circRNA varied notably between methods, the reported direction of regulation was largely consistent and overall remained concordant with the defined CIRI-DE baseline.

### 2.7 Inter-method concordance across DEA methods

Beyond the performance of DEA metho6ds on the tested scenarios, we next examined the concordance between the circRNAs identified by CIRI-DE and those detected by the other DEA pipelines when including information derived from the linear sources. Specifically, we collected circRNAs identified as significantly DE by both FSJ- and BSJ-only versions of all algorithms at three FDR thresholds (0.01, 0.05, and 0.10). This resulted in a set of significant circRNAs which served as the basis to assess the degree of overlap in the reported circRNAs between methods. We did not consider the directionality of expression at this stage. To maintain consistency for the previously reported results in the study, we explored the results obtained at an FDR < 0.05.

Initially, in the BC dataset from a total of 20 reported circRNAs, there were six jointly identified by both methods and only 3 were shared with the limma-voom methods. However, it is important to highlight that all circRNAs detected by edgeR-QLF were present within the broader set identified by CIRI-DE, which consistently reported a higher number of circRNAs as opposed to the rest of the methods.

Given the relatively low number of significantly DE circRNAs detected in the BC dataset, we next assessed whether this pattern continued in a higher signal dataset, such as EBC1, where a total of 98 significantly DE circRNAs were identified. (Figure 5C). From the results obtained, we could identify more than 30 circRNAs overlapping between CIRI-DE and the edgeR-QLF pipeline, as opposed to limma-voom and DESeq2, which reported more than 20 and 5 unique circRNA, respectively, not detected by the other pipelines. By contrast, the number of circRNAs jointly identified by DESeq2, edgeR-QLF, and CIRI-DE was only five.

This trend persisted across the datasets when evaluating concordance at FDR thresholds of 0.01 and 0.10 (Supplementary Figures 12 to 16).

### 2.8 Run time analysis

The computational efficiency of the benchmarked DEA tools was assessed by recording the wall-clock time required to complete end-to-end analysis across all 1140 simulated datasets. Methods were run a total of 13680 times, with 6840 runs each on the “ZeroSets” and “DE-Signal” datasets (Figure 6A). Total runtime varied considerably between methods, ranging from 1.4 minutes for edgeR to 15.2 minutes for limma, and 72.17 minutes for DESeq2 (Figure 6B).

**Figure 6.**
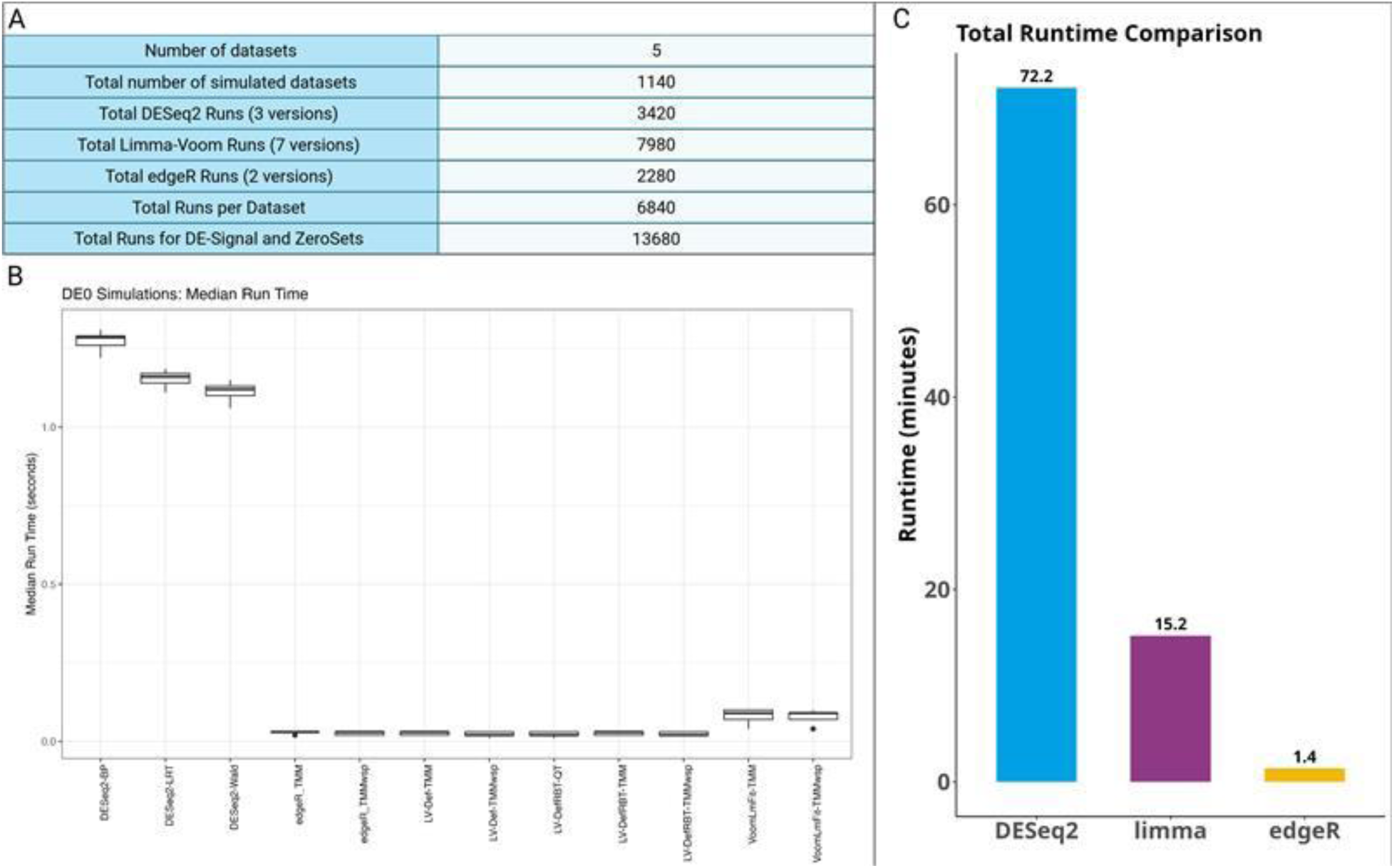
Run time statistics (A) Summary of total runs. (B) Median run time per tool (C) Comparison of time taken to run all DEA tools on the datasets.

At an individual dataset level such as the Zero Sets dataset, all tested methods demonstrated rapid performance, completing analysis within seconds. Median runtimes ranged from 0.01 seconds for edgeR to 1.37 seconds for DESeq2 (Figure 6C). This runtime pattern remained consistent across all the evaluated filtering approaches.

## 3 Discussion

In this two-part benchmarking study, we systematically evaluated the performance of four well-established DEA methods in circRNA data. In the first part, we assessed the effect of three different pre-filtering criteria on twelve different configurations of edgeR, DESeq2, and limma-voom, using simulated semi-parametric datasets. In the second part, we examined the impact of incorporating linear RNA and FSJ information into circRNA DEA, an approach that remains widely underexplored in the field.

Benchmarking was performed across five datasets, among them two in-house platelet-derived ones from healthy donors and early breast cancer patients. We selected platelets for their naturally elevated circRNA content compared to nucleated cells with reported circRNA enrichment levels ranging from 14- to 26-folds higher relative to RNAse R-treated samples (Alhasan et al., 2016). This unique characteristic of platelets enabled us to evaluate the performance of DEA methods in a physiologically circRNA-rich system and simultaneously avoid any potential bias introduced by exonuclease treatment in the abundance of linear RNA transcripts, which could have hindered the results of the second part of this benchmark.

Consistent with previous reports, both utilized detection algorithms identified a high number of circRNAs while also reporting a high percentage of zero counts across all datasets (Buratin et al., 2023). Despite the absence of circRNA enrichment, the platelet-derived datasets (EBC1 and EBC2) contained the highest number of circRNA post-filtering, further establishing platelets as a naturally enriched source of circRNAs, even in biological conditions such as early breast cancer where transcriptomic changes might not be as elevated as in late-stage breast cancer (Luo et al., 2020), reinforcing their potential as a suitable input material for biomarker detection without the need for additional enzymatic enrichment steps.

Moreover, the abundance of zero- and low-count circRNAs resulted in reduced inter-tool concordance as reflected in the low Jaccard similarity indices, which only improved following strict filtering steps. These properties of circRNA data reinforce the necessity to carefully define filtering thresholds prior to DEA. Moreover, filtering low-count transcripts is an essential preprocessing step in both bulk and single-cell RNA-sequencing workflows, where it has been consistently shown to improve the statistical power of DEA methods (Deyneko et al., 2022; Moutsopoulos et al., 2021; Sha et al., 2015). In sparse, zero-inflated circRNA count matrices, the importance of filtering becomes even more critical as it directly affects the ability to distinguish between biological signal and technical noise, algorithm-specific artifacts, or extremely low-abundance circRNA transcripts.

While the impact of different filtering strategies has already been explored in single-cell sequencing data (Gagnon et al., 2022; Soneson & Robinson, 2018a), its implications in circRNA data remain underexplored. Moreover, the filtering criteria applied to circRNA count matrices vary widely from study to study, reflecting a lack of consensus on what constitutes an appropriate threshold. We were presented with the same methodological uncertainty in our analysis, where the absence of standardized recommendations on the matter encouraged us to perform a systematic evaluation of different filtering strategies on the performance of widely utilized DEA methods.

The evaluated thresholds included the automated filterByExpr() approach, representing the strictest criterion, a minimum of more than 5 (Min 5) and 1 (Min 1) read counts, with the latter representing the most lenient approach explored. Although studies on low-count bulk sequencing data have suggested that careful model calibration alone may be sufficient to omit filtering altogether (Raithel et al., 2016), the applicability of these findings to circRNA data remains unclear, especially given the special characteristics of circRNA count matrices.

Our results indicate that auto-filtering represents the most stringent filtering strategy as reflected by the reduced zero inflation, higher median BSJ counts across all datasets, and increased concordance in the identified circRNAs between both detection algorithms. While Min 1 and Min 5 retained a considerably higher percentage of zero counts, this was at the expense of reduced concordance between CIRI3 and CE2, suggesting that these thresholds are not efficient in removing unreliably detected circRNAs or artifacts. Similar results were reported by Digby et al., 2023, when exploring the efficiency of a commonly used, albeit lenient, filtering step (≥ 2 reads at the BSJ site) in circRNA data. In sum, these findings suggest that auto-filtering represents a robust strategy for increasing confidence in circRNA candidates identified across different detection algorithms; however, additional research combining the results of more algorithms such as CircCompara (Gaffo et al., 2022), KNIFE (Szabo et al., 2015), circRNA_finder (Westholm et al., 2014) or find_circ (Memczak et al., 2013) is required to fully assess its robustness.

Furthermore, when evaluating the impact of the different filtering thresholds on the performance of the DEA methods, we found that most methods achieved lower FPR and FDR, alongside high sensitivity and high F1 scores on auto-filtered count matrices; a stark contrast compared to Min 1 filtering, in which methods suffered a notable decrease in performance. This drastic decline may be partly explained by the substantially higher zero count sparsity retained in Min 1 count matrices, which contained on average 5.7-fold more zero counts compared to the auto-filtered ones.

Moreover, we found that the applied filtering strategy largely impacted the ability of DEA methods to effectively control Type 1 error across all datasets. This effect was particularly evident in DESeq2 and edgeR, which maintained near nominal FPR values both under auto-filtering and Min 5 but exhibited conservative behavior when applied to Min 1-filtered matrices containing up to 28% zero counts. We did not observe any substantial performance differences across the tested DESeq2 model specifications.

As previously reported in RNA-sequencing data, these results suggest that retaining weakly supported BSJ counts influences dispersion estimation within the negative binomial models, ultimately reducing their statistical power to identify truly DE circRNAs (Berge et al., 2018).

These results are consistent with prior circRNA and other studies reporting the conservative behavior of DESeq2 with zero-inflated datasets (Abegaz et al., 2024; Q. Li et al., 2019; Buratin et al., 2023; Malhotra et al., 2021)

Interestingly, our findings suggest that limma-voom and, in particular, LV-Def and limma-voom-DefRBT provide a more balanced and stable trade-off between sensitivity and precision across different filtering conditions and heterogeneous circRNA datasets, derived from different biological sources, such as liquid biopsies and tissue, including here the particularly challenging HCC-PBMC dataset. This was consistently reflected by their overall stable performance under auto-filtering, displaying high F1 scores and AUPRC values. These results are in line with prior studies in circRNA and low-count mRNAs reporting on the stable performance of limma-voom in maintaing a balanced FDR control and sensitivity (Assefa et al., 2018; Buratin et al., 2023; Soneson & Delorenzi, 2013), establishing limma-voom as a reliable DEA tool for the analysis of circRNA data.

Another interesting observation was the notable decline in performance of DEA methods in the HCC-PBMC dataset, especially under lenient filtering thresholds. The poor performance on this dataset could be attributable to several factors, such as the naturally lower abundance of circRNAs compared to the nucleated blood cells and their biological composition and cellular heterogeneity (Nicolet et al., 2018). All these factors combined with the naturally low number of circRNAs may have resulted in a low-signal, high-variance scenario, which hindered the ability of DEA methods to distinguish noise from signal. These findings suggest the challenging nature of liquid biopsy-derived samples and raise concerns regarding the performance of DEA methods in such datasets. Nonetheless, circRNA enrichment in such scenarios can be an effective alternative to overcome such hurdles.

Given these limitations associated with heterogeneous datasets and challenging input materials, novel methods that account for the unique properties of circRNA can be beneficial to address these issues.

Several approaches have previously attempted to incorporate circRNA-specific information into DEA, including the use of linear-to-circular ratios at the BSJ sites such as DEBKS (Liu et al., 2022) and CircMeta (L. Chen et al., 2020), as well as the strategy implemented in CIRI-DE (Zheng et al., 2025), which normalized circRNA expression by borrowing information from FSJ reads at the corresponding loci.

Given that the CIRI-DE framework is currently among the more widely adopted approaches in literature, to the best of our knowledge, we provide the first independent benchmark evaluating whether the incorporation of FSJ-derived information improves circRNA detection compared to conventional DEA pipelines that omit such information.

Our results demonstrated an increase in the number of detected DE circRNAs when information from corresponding FSJs or linear transcripts was included alongside the BSJ counts. This suggests the potential presence of shared signal between circular and linear species that may be overlooked when circRNAs are analyzed in isolation using bulk RNA-seq frameworks. An interesting observation was the robustness of the glmLRT approach from edgeR, which is internally utilized by CIRI-DE, as it identified a higher number of DE circRNAs compared to the edgeR-glmQLFit pipeline. However, this difference may very well reflect the more stringent error control properties of the quasi-likelihood framework rather than the methodological superiority of CIRI-DE (Y. Chen et al., 2016, 2025a). Nonetheless, despite differences in the total number of identified circRNAs between both approaches, there was a substantial overlap between identified circRNAs.

Collectively, these findings suggest that incorporating linear transcript information prior to DEA can enhance circRNA detection, even when using statical models originally developed for bulk RNA-sequencing data. Furthermore, adopting stringent automated filtering is essential for bridging the gap between current bulk RNA-seq models and the specialized requirements of robust circRNA analysis, ultimately enhancing the reliability and clinical utility in non-invasive cancer diagnostics.

## 4 Conclusions

In conclusion, we provide a comprehensive evaluation of widely used differential expression methods when applied to non-circRNA-enriched RNA-sequencing libraries. Our analysis demonstrates that data preprocessing steps, particularly filtering strategies, greatly influence DEA method performance, with overly lenient thresholds substantially reducing their performance, with the exception of limma-voom methods which generally displayed a stable performance. While we strongly recommend dataset-driven filtering, we identified filterByExpr()-based automated filtering as an overall reliable alternative to arbitrary thresholds. Furthermore, we show that incorporating linear transcript information improves circRNA detection in non-enriched datasets; however, the applicability of this approach to RNAse R-enriched libraries requires further investigation. Overall, our findings highlight some of the limitations of utilizing RNA-sequencing DEA methods to circRNA data and support the further development of novel circRNA-specific models that consider sparsity and distributional properties and incorporate linear-derived information as well.

## 5 Methods

### 5.1 DEA method literature evaluation

A literature search was conducted to identify the most prevalent DEA methods applied in circRNA studies published in peer-reviewed journals between 2015 and 2025. Searches were performed in Google Scholar using the keywords “circRNA”, “expression”, and/or “differential expression”. Review articles, PhD dissertations, and conference abstracts were excluded. For each eligible publication, information on the DEA method(s) was manually extracted, including studies reporting on the use of multiple methods or combinations of analytical frameworks. In the end, a total of 100 research articles meeting the inclusion criteria were compiled for comparative assessment. Given the high frequency of combinatory approaches, unique combinations were not reported separately but rather collapsed into the “Others” category.

### 5.2 Online dataset selection

We selected three publicly available datasets comprising a total of 84 samples derived from two conditions: healthy controls and cancer samples. In two datasets, BC and HCC-Tissue, RNA was isolated from tumor tissues and matched adjacent normal tissues, whereas the third dataset consisted of PBMC samples from patients with hepatocellular carcinoma (HCC). Datasets were selected based on a listed set of criteria: (i) a minimum of five biological replicates per condition; (ii) a minimum sequencing depth of 20 million reads per sample; (iii) availability of raw sequencing data accompanied by comprehensive sample metadata, and (iv) clear separation at circRNA level between samples within each condition.

Raw sequencing data for each dataset were retrieved from the SRA public repository (Katz et al., 2022) and complemented with detailed sample metadata from the European Nucleotide Archive (Burgin et al., 2023). Initial quality assessment of the raw reads was performed using FastQC (Andrews, 2010), followed by preprocessing with fastp (S. Chen et al., 2018) using default parameters. Cleaned reads were then aligned to the human reference genome (Ensembl release 112) using BWA-MEM (H. Li, 2013) and HISAT2 (Kim et al., 2019) for CIRI3 (Zheng et al., 2025) and CircExplorer3 (Ma et al., 2019) respectively. For the quantification of linear mRNAs, the splice-aware aligner STAR (Dobin et al., 2013) was used. In those cases where read quantification was not integrated into the analysis pipeline, featureCounts (Liao et al., 2014) was utilized to quantify read counts for each genomic feature.

### 5.3 Platelet RNA library preparation and sequencing

We analyzed two in-house datasets to further evaluate the performance of DEA methods. Early-(stage) Breast Cancer 1 (EBC1) and Early-(stage) Breast Cancer 2 (EBC2) comprise 11 healthy donors and 9 patients from Békés County Central Hospital, Pándy Kálmán Member Hospital using protocol approved by the Hungarian Health Science Council Scientific and Research Ethics Committee Board. Blood was drawn into 9-mL ACD-A Vacuette tubes (Greiner Bio-One International GmbH) and processed within 36 h of venipuncture.

Platelet isolation and RNA extraction were performed according to the ThromboSeq protocol described by Best et al., 2019. Briefly, platelets were enriched by sequential centrifugation; the resulting platelet was resuspended in RNAlater (Thermo Fisher Scientific) and incubated overnight at 4 °C.

Total RNA was purified using the miRNeasy Micro Kit (Qiagen) according to the manufacturer’s instructions. RNA libraries were prepared using the NEBNext® Ultra™ II Directional RNA Library Prep Kit for Illumina® (New England Biolabs).

Sequencing was performed on an Illumina NextSeq 2000 platform generating paired-end 150-bp reads. Raw files were processed as previously described in Section 5.2.

### 5.4 Statistical analysis

Inter-sample similarity within each condition was assessed by calculating Spearman correlation coefficients using the stats R package (R Core Team, 2025). To evaluate the concordance between CIRI3- and CE2-retained circRNAs following filtering, the overlap was quantified using the Jaccard similarity index, computed with the *Jaccard* R package (version 0.1.2) (Chung et al., 2019).

To assess differences in circRNA expressions between conditions, PCA was performed to test three different scenarios: (i) using BSJ counts alone, (ii) FSJ counts alone, (iii) a combination of both BSJ and FSJ counts. For each scenario, the corresponding count matrices were first filtered using the filterByExpr() function from edgeR (Y. Chen et al., 2025a) to remove zero and lowly expressed counts. DGEList objects were then constructed separately. For the approach implemented in CIRI3, normalization factors were estimated from FSJ counts and then subsequently applied to the DGEList objects constructed from BSJ counts. This strategy integrates linear RNA information to stabilize normalization when modeling circRNA abundance. Read counts were normalized using the TMM method implemented in the edgeR package, due to its robustness to outlier samples.

### 5.5 Simulation of datasets

Using the selected publicly available datasets along with our in-house platelet datasets, we generated two types of *in silico* datasets to benchmark DEA tool performance; a “zero set” dataset containing no DE circRNAs was generated to assess their Type I error control, and datasets containing 10% DE circRNAs to evaluate their performance using different metrics that require the ground truth set. All *in silico* datasets were generated using SPsimSeq, a semi-parametric simulation framework that preserves key characteristics of the input data, including mean expression distributions, sample variability, as well as the high proportion of zero counts, typical of circRNA expression data (Assefa et al., 2020). Prior to the generation step, the original BSJ feature count matrices were filtered in three different scenarios: (1) automatic filtering using edgeR’s “filterByExpr()” with default parameters, (2) with min.count set to 1, and (3) min.count set to 5.

Next, due to substantial variations in BSJ counts across samples within each condition, the IQR outlier detection method was applied separately to each group to identify both upper and lower outlier samples, reducing potential bias and ensuring a controlled intra-condition variability. This resulted in the removal of one breast cancer tissue sample from the BC dataset (5 BC samples vs 6 healthy controls) and one sample from the EBC2 dataset, resulting in five healthy and four patient samples. No outlier samples were detected in the EBC1 dataset; therefore, all ten samples were utilized for the simulation analysis. Additionally, two HCC samples were removed from the HCC-Tissue dataset (17 healthy and 15 HCC), and three healthy samples were removed from the HCC-PBMC dataset, resulting in 15 healthy and 18 cancer samples. Depending on the size of the original dataset as determined by the sample number per condition, we performed 30 (BC, EBC1, and EBC2) and 50 simulations (HCC-PBMC and HCC-Tissue), modeling a minimum of 1,000 genes. Library sizes across all simulations were kept consistently the same.

For the 10% or Signal-10 dataset, DE circRNAs between the two conditions were defined by an absolute log fold change ≥0.5. All the simulated datasets were then post-processed by initially inspecting their quality and distribution patterns and finally filtered to remove zero counts. Quality of the simulated datasets was inspected using countsimQC (Soneson & Robinson, 2018b).

We retained the original study design to ensure that the algorithms could distinguish between the two conditions present in the data. Each simulation modeled 1,000 circRNAs, including a null scenario with no DE circRNAs and a differential scenario in which 10% of the circRNAs were DE. The two conditions were simulated in balanced proportions, with no batch effects introduced. All remaining parameters were kept in their default state.

### 5.6 Selection of DEA methods

To evaluate the BSJ-only and combined FSJ-BSJ approaches, we assessed the performance of four widely utilized DEA methods identified through edgeR, limma-voom (Ritchie et al., 2015), DESeq2 (Love et al., 2014), and CIRI-DE (Zheng et al., 2025). Though most surveyed studies utilized these methods using default parameters, we evaluated their performance both at the default as well as adjusted parameter settings to account for the zero-heavy nature of circRNA data.

Specifically, for DESeq2, we tested the Likelihood Ratio Test (LRT), Wald Test, and betaPrior = TRUE, whereas for edgeR we applied the quasi-likelihood F-test with both TMM and TMM with singleton pairings (wsp) normalization, with the latter performing better on sparse data. For limma-voom, we tested and evaluated three main configurations, including the standard pipeline (voom:lmFit:eBayes), LmFit with robust = TRUE, and voomLmFit (sample.weights = TRUE). All these methods were tested on count matrices normalized in two ways, with TMM and TMMwsp. In total, this resulted in 14 different method variants.

### 5.7 Evaluating the impact of linear RNA normalization in circRNA differential expression

To evaluate the effect of employing gene-level normalization from predicted FSJs, we extended the analysis to include the CIRIquant framework, which normalizes circRNA counts using gene-level information derived from linear transcript expression (Zheng et al., 2025). This approach entails constructing a DGEList object from the filtered and normalized FSJ count matrices produced by CIRI3 during the circRNA identification pipeline and applying these post-filtered FSJ library sizes along with gene-level normalization factors to the corresponding BSJ counts.

DESeq2 does not natively support edgeR’s calcNormFactors function for normalization, hence preventing direct application of the CIRIquant strategy of FSJ-derived normalization factors. To implement an equivalent approach, we derived size factors from DESeq2’s median-of-ratios method and applied them to the BSJ matrix. Following this external normalization, the standard DESeq2 dispersion estimation and DEA testing were performed with various settings.

In addition to the FSJ-based normalization, we assessed the effect of using total linear RNA count output from STAR. This step was performed by filtering the counts using edgeR’s filterByExpr() with default parameters, followed by TMM or TMMwsp normalization and DGEList construction. The filtered and normalized library sizes were then applied to the BSJ matrices as previously described. For this analysis, we surpassed the original glmFit model implemented in the CIRI-quant framework by also evaluating the quasi-likelihood F-test, which is also the currently recommended default for edgeR workflows (Y. Chen et al., 2025b). This resulted in six configurations listed as follows: (1) CIRI-quant-BSJ-glmFit, (2) CIRI-quant-FSJ+BSJ-glmFit, (3) CIRI-quant-BSJ-qlmFit, (4) CIRI-quant-BSJ+FSJ-qlmFit, (5) STAR-CIRI-BSJ-glmFit, and (6) STAR-CIRI-FSJ+BSJ-glmFit.

These analyses were carried out on five experimental datasets that served as the starting point for the simulation analysis in the first part of the study. Each dataset was analyzed using a standardized preprocessing pipeline, starting with filtering using filterByExpr(), followed by normalization with TMM or TMMwsp, and lastly DGEList construction. For DESeq2, the internal median-of-ratios normalization was left unchanged. The linear normalized circRNA counts were then used for DEA to identify circRNAs between the tested conditions.

As no ground truth DE set was available for the experimental datasets, we focused primarily on comparing the total number of significantly differentially expressed circRNAs (FDR < 0.05), the amount of overlap of the significantly expressed detected circRNAs across methods, and the distribution of the p-values.

### 5.8 Type 1 error control

To evaluate Type I error control, we generated null datasets (ZeroSets) with no true DE between the tested conditions. Under the null hypothesis, any circRNA identified as significant represents a false positive; therefore, FPR was calculated as the proportion of false positives over the total number of tested circRNAs. For each DEA method, we calculated the FPR as the proportion of circRNAs with unadjusted p-values below three nominal thresholds: 0.01, 0.05, and 0.10. The same analysis was performed to calculate FPR values across three different filtering strategies.

### 5.9 Performance evaluation on the DE-Signal dataset

For each differential expression method, we computed TPR, FDR, F1 Score, and AUPRC across 195 simulated datasets. These calculations were performed at three significance levels (α ∈ {0.01, 0.05, 0.10} and filtering strategies (detailed in Methods Section 5.5). Per method, medians of all metrics were calculated and reported across datasets to quantify their central tendency and per-dataset variability. All analyses were performed in R version 4.5.0 using custom scripts, whereas the AUPRC analysis was carried out using the PRROC package Version 1.4 (Grau et al., 2015).

The consistency and reproducibility of differential expression calls across simulated replicates per dataset was examined by quantifying the overlap of significantly expressed circRNAs within each dataset using the Jaccard similarity index (Jaccard, 1912). Jaccard index measures the similarity between two sets of data as the proportion of shared elements relative to the total number of unique elements present in either set, thus making it a robust metric to differences in dataset size. Values range from 0 (methods disagree on circRNAs identified as DE) to 1 (methods consistently detect the same circRNAs as DE), with higher values indicating greater agreement between methods. To calculate the Jaccard index, the result files for each replicate were parsed to retain circRNAs with FDR ≤ 0.05. Replicates were then compared pairwise within each dataset by counting the number of shared circRNAs between them and the number of unique circRNAs identified in either replicate. The obtained values were averaged per file, excluding the diagonals, which always result in a Jaccard value of 1 due to self-comparison.

### 5.10 Run time analysis

Computational efficiency for all the tested DEA tools was assessed by measuring the wall-clock time for complete end-to-end DEA analysis pipelines using R’s Sys.time() function. Runtime was recorded for all methods on both ZeroSets and DE-Signal simulated datasets. All analysis were performed on a workstation with the following specifications: [AMD Ryzen^TM^ 9 9950X processor, 128 GB RAM, running Ubuntu 25.04, R version 4.5.0].

## 6 Declarations

### 6.1 Availability of data and materials

The RNA-sequencing data generated in this study are available at NCBI SRA under BioProject ID PRJNA1429817. The code utilized to analyze the data in this study are available in the GitHub repository based on the following link: (https://github.com/Tin-26/circRNA-DE-method-evaluation).

### 6.2 Competing interests

The authors declare that they have no competing interests.

### 6.3 Funding

This project received funding from the National Research, Development, and Innovation Office (2023-1.1.1-PIACI_FÓKUSZ-2024-00029; 2024-1.1.1-KKV_FÓKUSZ-2024-00019; TKP-31-8/PALY-2021, 2025-1.1.2-GYORSÍTÓSÁV-2025-00038).

### 6.4 Authors’ contributions

E.Q. and V.V. contributed equally to this work. E.Q., V.V., K.P. conceived of the study design, and K.P. implemented the liquid biopsy experiments. K.Z. and K.P. conceptualized the patient cohorts. D.L. performed experimental laboratory work. D.L and B.C. coordinated sequencing of the Illumina RNA-seq samples. E.Q. wrote the analysis code. E.Q. and V.V. performed formal analysis. E.P., E.Q., V.V. and B.T. created visualizations and drafted the manuscript. G.J. made the code review. L.P. contributed to conceptualization and resources. L.T. and A.B. provided patient samples and clinical data. E.Q., B.T. and V.V. analyzed the results. E.Q. and V.V. drafted the original manuscript. L.H. supervised the work. All authors reviewed, edited, and approved of the final manuscript.

## 6.5 Acknowledgements

We thank study nurses at the Pándy Kálmán Member Hospital for managing sample collection. We acknowledge the institutional Sequencing Core Facility for sequencing support. We are particularly grateful to Gabriella Tick for editorial assistance and language proofreading.

## 6.6 Ethics approval and consent to participate

The Hungarian Health Science Council Scientific and Research Ethics Committee Board (ETT-TUKEB) approved this study (BM/32453-3/2024), and our research complies with all relevant ethical regulations. All participants gave their written informed consent.

## 6.7 Consent for publication

Not applicable.

## 7 Supplementary Materials

Additional file 1: Supplementary Figures. Supplementary Figures S1-S16.

Additional file 2: Supplementary Tables. Supplementary Tables S1-S4.

## Notes

### Competing Interest Statement

The authors have declared no competing interest.

